# Characterizing variants of unknown significance in rhodopsin: a functional genomics approach

**DOI:** 10.1101/512897

**Authors:** Aliete Wan, Emily Place, Eric A. Pierce, Jason Comander

## Abstract

Characterizing the pathogenicity of DNA sequence variants of unknown significance (VUS) is a major bottleneck in human genetics, and is increasingly important in determining which patients with inherited retinal diseases could benefit from gene therapy. A library of 210 rhodopsin (*RHO*) variants from literature and in-house genetic diagnostic testing was created to efficiently detect pathogenic *RHO* variants that fail to express on the cell surface. This study, while focused on *RHO*, demonstrates a streamlined, generalizable method for detecting pathogenic VUS. A relatively simple next generation sequencing (NGS)-based readout was developed so that a flow cytometry-based assay could be performed simultaneously on all variants in a pooled format, without the need for barcodes or viral transduction. The resulting dataset characterized surface expression of every *RHO* library variant with a high degree of reproducibility (Z’=0.94, R^2^=0.92-0.95), recategorizing 37 variants. For example, three retinitis pigmentosa pedigrees were solved by identifying VUS which showed low expression levels (G18D, G101V, P180T). Results were validated across multiple assays and correlated with clinical disease severity. This study presents a parallelized, higher-throughput cell-based assay for the functional characterization of VUS in rhodopsin, and can be applied more broadly to other inherited retinal disease genes and other disorders.

## Introduction

With the increasing availability and use of human DNA sequencing, the problem of accurately characterizing newly-discovered DNA variants has become a major issue in human genetics (Lappalainen, 2015; Richards, 2015). Difficulties in predicting the pathogenicity of DNA “variants of unknown significance” (VUS), even with all available bioinformatic, functional, and human data, routinely produce an ambiguous final result of genetic testing (Davies, 2012; Bean, 2013; Richards, 2015). Dealing with this ambiguity is a major problem for medical geneticists and genetic counselors who have to manage this uncertainty with patients who are expecting clarity. For example, grading systems have been developed to rank each variant on a scale from one to five, representing “pathogenic”, “likely pathogenic”, “uncertain significance”, “likely benign”, and “benign”. Evidence supporting pathogenicity is divided into additional categories, and counting the number of evidence points from each evidence category results in a final rank (Richards, 2015). Such complex systems can attempt to manage uncertainty in pathogenicity, but clearly it would be preferable to have more information about a variant to decrease the uncertainty level.

Current bioinformatic approaches to predicting variant pathogenicity are not sufficient to avoid these problems (Richards, 2015). For example, most algorithms for predicting the pathogenicity of missense mutations are only 65-80% accurate,(Thusberg, 2011) and specificity for the predictions of pathogenicity of missense and splice variants can be low (Vreeswijk, 2009; Choi, 2012; Houdayer, 2012). The latest techniques are raising the quality of the predictions(Gray, 2018), but in the end, the state-of-the-art is that “it is not recommended that these [bioinformatic] predictions be used as a sole source of evidence to make a clinical assertion” about a potentially pathogenic variant (Richards, 2015). In fact, “well-established *functional* studies showing a deleterious effect” is considered two levels-of-evidence higher than “multiple lines of *computational* evidence support[ing] a deleterious effect on the gene/gene product” (italics added)(Richards, 2015).

VUS not only cloud the interpretation and utility of clinical diagnostic testing, but also can lead to outliers and ambiguities when analyzing structure-function relationships of proteins of interest (Rakoczy, 2011). For example, when correlating the computational prediction of misfolding propensity and the age of onset of disease among rhodopsin mutants, some mutations are considered outliers and excluded from the regression analyses(Rakoczy, 2011); however, it is not clear whether the outliers could be due to imperfect computational models or to miscategorization of the mutant. Thus, improving the characterization of DNA variants is scientific importance, and has been included U.S. federal research priorities and identified as a knowledge gap in the understanding of inherited retinal diseases (National Eye Institute, 2012; Duncan, 2018).

This study implements an improved method to characterize potentially pathogenic DNA variants causing retinitis pigmentosa (RP). RP accounts for up to 25% of blindness or visual impairment in working age people (21-60 years) (Hata, 2003; Buch, 2004; Al-Merjan, 2005; Hartong, 2006), and therefore is an important cause of vision loss. Although RP is a Mendelian disease, it is genetically very heterogeneous, with mutations in over 60 different genes that can cause nonsyndromic RP (Daiger, 1998; Berger, 2010; RetNet). Genetic testing to identify the cause of disease has become increasingly important as more clinical trials for RP focus on patient populations with specific genotypes, e.g. studies recruiting *MERTK-, MYO7A-, PDE6A-, PDE6B-, RPGR-, orRLBP1*-affected RP patients (U.S. National Institutes of Health, 2017), as well as the occasional RP patient due to RPE65 mutations who would be eligible for the first FDA-approved gene therapy, voretigene neparvovec (Luxturna) (Russell, 2017). Patients without a genetic diagnosis are not eligible for gene specific treatments. Despite the practical importance of obtaining a genetic diagnosis, definitive causal variant(s) can be found in only about half of patients, and slightly more using the latest next generation sequencing (NGS) techniques (Neveling, 2012; Corton, 2013; Glockle, 2014; Wang, 2014; Consugar, 2015; Huang, 2015). For this reason, improving the characterization of DNA variants in RP is also of practical importance.

This study focuses on rhodopsin (*RHO*) mutants because *RHO* has the largest set of known pathogenic variants of any dominant RP gene, and among those genes, the structural, biochemical, and cell biological understanding of *RHO* is unmatched (Dryja, 1990; Mendes, 2005; Mendes, 2008; Krebs, 2010; Mendes, 2010; Rakoczy, 2011; McKeone, 2014; Athanasiou, 2018; Behnen, 2018). The rich body of existing data provides a context for interpreting data on new variants, and conversely, provides an opportunity to refine existing models of *RHO* structure and function (Rakoczy, 2011). A recent review (Athanasiou, 2018) notes that ongoing questions about the pathogenicity of *RHO* variants “reinforce the need for thorough genetics, such as segregation analyses, and in-depth functional analyses to confirm pathogenicity.” (Note that the term “variant” is used in this study to include any sequence change, whether that sequence change is a “mutant” that is known to be pathogenic, a VUS, or a benign change such as a synonymous control.)

Standard experimental methods of assaying *RHO* variants include assessment of surface expression / subcellular localization in cell-based assays (Sung, 1993; Li, 1998; Chuang, 2004; Chen, 2011; Toledo, 2011; Davies, 2012; Hollingsworth, 2013; Liu, 2013; McKeone, 2014; Yamasaki, 2014; Behnen, 2018) and assessment of bulk biochemical properties (Bosch, 2003; Dizhoor, 2008; Krebs, 2010; Bosch-Presegue, 2011; Opefi, 2013). These assays are performed on one variant at a time, and, especially for the biochemical assays, do not scale easily for larger numbers of variants.

The purpose of this study was to develop higher-throughput cell-biological methods to evaluate *RHO* variants for pathogenicity e.g. a “functional genomics” approach. We hypothesized that this approach would result in improved categorization of *RHO* DNA variants compared to using computational information alone. We also hypothesized that pedigrees with inconclusive genetic testing results and a *RHO* VUS could be solved using the functional data. This study presents a streamlined functional genomic screen (Figure 1) suitable for assaying hundreds of variants, using standard mutagenesis, transfection, flow cytometry, and NGS amplicon sequencing techniques that are broadly available. The bioinformatic analysis was custom-built, but also straightforward. More intricate functional genomic screens have been recently developed that interrogate thousands of variants at once(Melnikov, 2014; Brenan, 2016; Gasperini, 2016; Findlay, 2018) but these methods require specialized library construction techniques, lentivirus packaging, and viral transduction followed by selection, which were not needed for this study.

**Figure 1.**
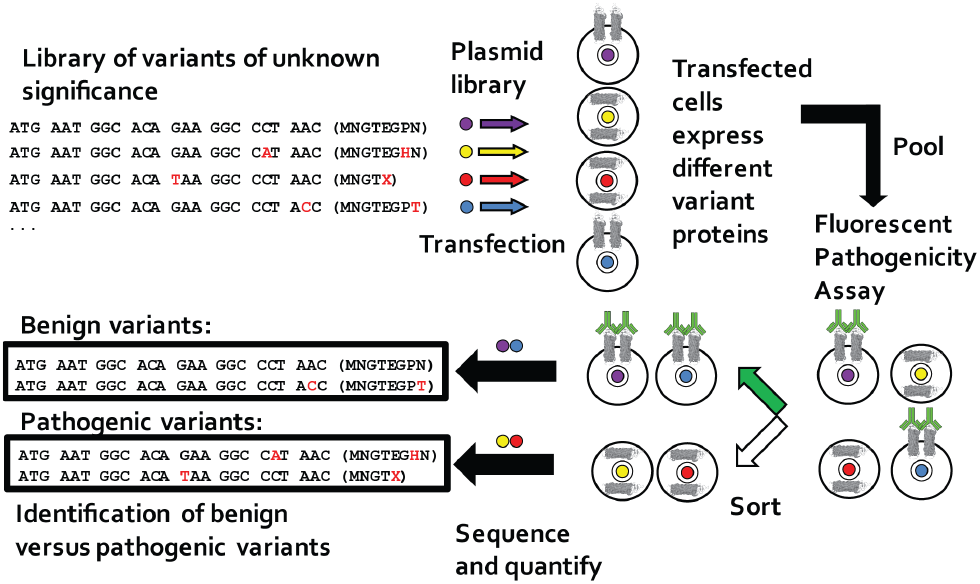
Experimental design of library screening for VUS characterization. An expression plasmid library of *RHO* variants was constructed using mutagenesis. (Four variants are shown as an example.) Each variant plasmid was transfected into cells, which were then pooled after transfection. The cells, after expressing the desired variant (grey), were stained using a fluorescent pathogenicity assay based on RHO surface expression (green). Fluorescence activated cell sorting (FACS) was used to separate high- and low-expressing cells. Each transfected cell carries with it the DNA of the variant of interest, and this “tag” can be sequenced using NGS and therefore count the relative frequency of each DNA variant in each pool, quantifying the pathogenicity of each variant in parallel.

This functional genomics approach was used to create a library of *RHO* variants and precisely assay their surface expression in an efficient, pooled format.

## Materials and Methods

### Identification of rhodopsin variants

A literature review (PubMed) and database search (HGMD, ClinVar) identified 211 known rhodopsin variants. Twenty-two database variants that were large insertions, large deletions, high-allele-frequency synonymous alleles, synthetic, or intronic (since the cDNA template has no introns) were not constructed. See Table S1. The remaining 189 variants were added to the variant library and of those, 175 were known as pathogenic, causing dominant RP, recessive RP, or congenital stationary night blindness. (Table S1) Fourteen variants were of uncertain significance or thought to be benign polymorphisms. All of the variants were well distributed across the cDNA, except that there were no variants between bp 664 and 745.

Eleven synonymous mutations, expected to have similar expression to wildtype and to serve as controls, were designed synthetically. The mutations were placed near the 3’ end of the gene in order to minimize any possible (though unexpected) changes to transcriptional or translational processes (Stoletzki, 2007). In addition, a search for rare (detected <3 times in our internal database, ExAC allele frequency<10^−4^) rhodopsin variants of unknown significance (VUS) from patient DNA sequencing performed in our institution’s genetic testing service (Consugar, 2015), identified 10 variants spanning 11 probands. All of these variants were novel (previously unreported) at the beginning of this study. Altogether, 232 rhodopsin variants were identified and 210 were ultimately constructed as a variant library: 189 variants from literature / databases, 11 synonymous controls, and 10 novel VUS.

The study protocol adhered to the tenets of the Declaration of Helsinki and was approved by the Institutional Review Board of Massachusetts Eye and Ear.

### Rhodopsin variant cDNA library construction

A Gateway destination vector was created that contained: a CMV early enhancer/chicken β actin (CAG) promoter, V5 tag, Gateway cassette (Life Technologies), internal ribosome entry site (IRES), mCherry fluorescent protein marker (Addgene), and ampicillin resistance gene. Plasmids were propagated in ccdb survival cells (Life Technologies) with chloramphenicol and ampicillin. A pDONR entry clone containing the human rhodopsin cDNA sequence was purchased (GeneCopoeia #GC-T1321, Rockville, MD). Each of the 210 *RHO* variants was created via site-directed mutagenesis using a one-primer modification to the QuikChange II protocol from Agilent.((Braman, 1996) and (http://qb3.berkeley.edu/macrolab/quick-change-mutagenesis/). Primers were synthesized in 96-well plates by Integrated DNA Technologies (Coralville, Iowa). Entry clone plasmids were propagated in One Shot Top10 chemically competent E.coli (Invitrogen) with kanamycin 50 micrograms/mL. The 1047 bp rhodopsin insert and flanking att sites were Sanger sequenced in both directions to exclude second-site mutations. Reracked positive clones were re-sequenced to verify the identity of each clone on each plate.

Expression clones (N=210) were created by recombination of destination vector and each of the mutagenized entry clones, using LR Clonase II (Invitrogen). The resulting expression clone consisted of: pCAG-V5-*RHO*(*WT or variant*)-IRES-mCherry and was propagated in Stbl3 cells (Invitrogen) with ampicillin 50-100 micrograms/mL. Clones were sequenced, reracked, and resequenced to verify the identity of each clone on each final library plate. DNA purification and sequencing (96-well plate) was performed by the CCIB DNA Core Facility at Massachusetts General Hospital (Cambridge, MA).

### Cell Culture

HEK293 cells (ATCC) were cultured using aseptic technique and grown in 10% fetal bovine serum (FBS) in Dulbecco’s modified Eagle serum (DMEM) without antibiotics. Cells were grown at 37°C and 5% CO_2_ in a standard cell culture incubator and passaged at subconfluent densities every 2-3 days as needed.

### *RHO* transfection and surface expression assays by immunofluorescence

All transfections were performed using Lipofectamine 2000 (ThermoFisher 11668019) on HEK293 cells seeded at a density of 200 cells/mm^2^ on tissue culture dishes. The day after seeding, the media were refreshed and the transfection mixture added as described below. Samples were collected 48 hours after transfection.

#### Immunofluorescence microscopy

Glass coverslips were placed in wells before cell seeding into 6-well dishes. Transfection was performed using 2 μg plasmid DNA and 3 μL Lipofectamine in 150 μL Opti-MEM I (ThermoFisher 31985062). At collection, cells on coverslips were fixed in 4% PFA for 20 min and blocked with 3% BSA in PBS for 10 min before applying Ret-P1 anti-rhodopsin antibody (Sigma #O4886) at a final concentration of 1:1000 in blocking buffer for 1 hr. After washing with PBS, Alexafluor-488 goat anti-mouse antibody (ThermoFisher A-11029) was applied at a final concentration of 1:300 in blocking buffer for 1 hr and Hoechst 33342 (ThermoFisher #H3570) applied for 1-5 min at a concentration of 1:5000. Coverslips were mounted onto slides with Fluoromount G and dried overnight at room temperature. Slides were kept at 4C before viewing on a Nikon TI Eclipse microscope or Leica TCS SP5 confocal microscope.

#### Flow cytometry

Transfection was performed in 24-well plates using 0.5 μg plasmid and 1.5 μL Lipofectamine in 50 μL Opti-MEM I. At collection, the cells were washed briefly with PBS and then 200 μL trypsin added and allowed to incubate for 4 min at 37°C. A 300 μL aliquot full media was then added, and cells collected into 2.2 mL tubes. The cells were centrifuged at 400g in a floor centrifuge for 4 min and the supernatant aspirated. Cells were fixed in 4% PFA for 20 min and blocked with 3% BSA in PBS before applying the Ret-P1 primary antibody at a final concentration of 1:1000 in blocking buffer for 30 min. After washing with PBS, Alexfluor-488 secondary antibody was applied at a final concentration of 1:300 in blocking buffer for 30min. After a final PBS wash, cells were analyzed on a BD LSRII Flow Cytometer.

Flow cytometry results were analyzed with Flowing Software (http://www.flowingsoftware.com). The percentage of cells with high RHO surface expression was determined by setting the quadrants based on the WT and P23H controls in each experiment and then dividing the percent of cells in the top right quadrant (double-positives) by the sum of the top right and bottom right quadrants (all transfected cells). Resulting data were averaged over independent transfections from different weeks (N=1-3).

#### Fluorescence-activated cell sorting (FACS)

Cells were prepared as described for flow cytometry analysis but pooled after transfection and then fixed in zinc-based fixative (ZBF) (BD Pharmingen #550523) instead of 4% PFA. (ZBF was used because PFA fixation/crosslinking prevented downstream PCR amplification of residual plasmid DNA(Wester, 2003), while unfixed cells did not stain with Ret-P1 antibody.) All subsequent washes substituted TBS (tris buffered saline) for PBS to reduce salt precipitation. Cells were sorted into TBS or PBS with 1% FBS using a BD SORP 5 Laser Vantage SE DIVA. Replicate samples were derived from independent transfections on different weeks (N=3).

Additional experiments to implement a pooled transfection method were not effective; at standard plasmid concentrations, large numbers of different variants entered each cell causing every cell to stain similarly, while at low plasmid concentrations, not enough RHO expression was achieved for robust staining and cell sorting (not shown).

### Rhodopsin cDNA or RNA extraction and amplification from sorted cells

Immediately after sorting, cells (from 4 × 10^4^ to 2.3 × 10^6^) were pelleted at 400xg for 4 min and resuspended in 700 μL RLT buffer (Qiagen) with beta-mercaptoethanol (1:100), mixed and aliquoted into two samples of 350 μL each and stored at −80° C until extraction. Plasmid DNA and total RNA were isolated. Preliminary experiments (not shown) demonstrated that RNEasy mini columns (Qiagen #74104) can isolate both RNA and the relatively low molecular weight plasmid DNA in one step using the manufacturer’s standard RNA isolation protocol. For total RNA extraction, the manufacturer’s standard RNEasy protocol was followed, including on-column DNAse digestion (Qiagen #79254). For plasmid DNA extraction, a separate aliquot of cells in RLT buffer was purified using RNEasy mini columns without DNAse digestion, and PCR performed without reverse transcription to avoid amplifying RNA. The final RNA or plasmid DNA/RNA mixture was eluted in 35 μL water, and 15 μL was used for the RT-PCR or PCR reaction described below.

For DNA-extracted samples, PCR amplification was performed using forward primer 5’-GTTTGTACAAAAAAGCAGG-3’ and reverse primer 5’-GGAATTTACGTAGCGGC-3,’ which were complementary to regions of the plasmid DNA flanking the rhodopsin sequence. HotStarTaq polymerase (Qiagen #203203) was used with the following PCR program: 1) 95°C for 15 min; 2) 36 cycles of: 94°C for 30 sec, 50°C for 30 sec, 72°C for 1 min; 3) 72°C for 10 min. For RNA-extracted samples, One-Step RT-PCR reagent (Qiagen # 210212) was used to create cDNA and then the cDNA was amplified by PCR. The following program was used: 1) 50°C for 30 min; 2) 95°C for 15 min; 3) 36 cycles 94°C for 30 sec, 50°C for 30 sec, 72°C for 1 min; 4) 72°C for 10 min. One μL of each 100 μL reaction was run on a Tapestation (Agilent) to determine approximate size and concentration and to confirm presence of a single band of amplified template. Negative control reactions (minus RT for RNA, plus DNAse for DNA) showed no cross-amplification between samples that were intended to amplify RNA or DNA, respectively. PCR reactions were processed through a PCR clean-up column (Qiagen #28104), and the concentration determined using a QuBit dsDNA HS Assay (ThermoFisher Q32854).

### NGS amplicon sequencing without library barcodes

Smaller scale NGS sequencing (1/96 of a MiSeq run per sample) was performed by the CCIB DNA Core Facility at Massachusetts General Hospital (Cambridge, MA). For more read depth (~1/10 of a MiSeq run per sample), NGS sequencing was performed in the Ocular Genomics Institute facility (https://oculargenomics.meei.harvard.edu/index.php/gc) using the following protocol: the PCR product was sheared on a Covaris E220 focused ultrasonicator set to a treatment time of 360 sec, acoustic duty factor of 10%, peak incident power of 175 W and 200 cycles per burst. Library preparation was performed with the Truseq nano LT kit (Illumina #FC-121-4001), but modified to use AMpure XP beads for the clean-up steps (Beckman Coulter #A63881). Briefly, steps included: cleanup, quantification, end repair, clean up, A-tail, adapter ligation, clean up, PCR enrichment, clean up, quantification and normalization, denature, and run on MiSeq using 2 × 121 cycles with a 6 bp index. Because of relative overrepresentation of the amplicon ends, PhiX library (10%) was added to the final mixture.

### Bioinformatic quantitation of low frequency variants in NGS amplicon sequencing

Most variant callers are designed to work with diploid genomes and do not call variants present at less than about 50% frequency. Extensive testing with specialized low-frequency variant callers (Spencer, 2014) showed that these variant callers, including “lofreq” (Wilm, 2012), did not accurately call all variants, particularly small insertions/deletions (a known limitation of alignment algorithms) (Jiang, 2015). Additional analyses with MuTect (Cibulskis, 2013) or MuTect2 showed low sensitivity using default settings, and settings with appropriate sensitivity and specificity were not identified(not shown). Multinucleotide polymorphisms (e.g. 511_512delCCinsGA) were particularly problematic.

Therefore, an alignment-free approach was used to detect and quantify low frequency variants in NGS data when the variants are known. This algorithm uses no alignment step and simply counts the number of exact-match occurrences of a 20 bp “probe” sequence string designed for each known variant. The “probes” (in analogy to Southern blot probes), were designed *in silico* containing each of the known 210 nucleotide changes. In most cases, the primer used for QuikChange mutagenesis was simply trimmed to 20 bp to create the “mutant probe”. A corresponding “wildtype probe” was created to quantify and normalize for variations in coverage. To estimate signal-to-noise ratio in the context of NGS sequencing errors(Fox, 2014), a “noise probe” was manually created containing an alternate nucleotide change which was not present in the wildtype or mutant probes. Each occurrence of these 630 probes (wildtype, mutant, or noise × 210 variants) was counted in the text of each sample’s FASTQ file using the Linux tool *grep*, similar to a strategy previously published (Bujakowska, 2015). The coverage at both ends of the amplicon was overrepresented, presumably due to sequencing of residual unsheared amplicon.

Preliminary experiments showed that the signal-to-noise ratio was low for some variants, particularly at amplicon ends, where early MiSeq cycles had a lower base call quality value. Thus, before quantifying the frequency of each probe in each FASTQ file, a script was implemented to censor all FASTQ files on a base pair-by-base pair basis for a high-quality Q score from 36-41 (“E” to “J”), resulting in improved signal-to-noise ratio (not shown). For graphing, signal-to-noise ratios were capped at 1000 when zero noise probes were detected. Run-time for quantification of the entire dataset was about 2 hours on our local computer cluster. These scripts are available at https://github.com/jcomand/VariantCounting.

The experimental design called for relative quantitation of each variant in two pools of transfected and sorted cells, in this case “high” versus “low” surface-RHO-expressing cells. The raw read counts of each mutant probe were normalized to the read depth of the wildtype probe. i.e. the final “NGS ratio” used for analysis was the number of read counts: (mutant probe_high/wildtype probe_high) / (mutant probe_low/wildtype probe_low). Three independent biological experiments (e.g. separate transfections on different weeks) were converted to log-ratios for calculation of averages and standard errors for each variant.

### Predicting clinical disease severity

Clinical outcomes of subjects with *RHO* mutations have previously been published, including by our institution(Berson, 2002). Electroretinography (ERG) data were analyzed from these subjects and subsequent subjects with *RHO* mutations from later years. The baseline (first visit) 30 Hz cone flicker ERG amplitude was used as the clinical outcome measure, as it has a broad dynamic range (>3 log units) across disease severities (Berson, 2002). Other ERG parameters or other outcomes such as visual field area and visual acuity were not used, to avoid multiple testing. To maximize sample size, data from DNA variants with the same amino acid change were pooled. Of the amino acid changes represented in the clinical data, class 2 (misfolding) mutations were the only biochemical category(Rakoczy, 2011) that had more than two mutations represented. The analysis was therefore limited to subjects with class 2 mutations (N=69 subjects). (Class 2 mutations also have the best biological rationale and precedent for correlating with disease severity in this assay (Rakoczy, 2011; Athanasiou, 2018).) Standard linear regression was used to predict the logged 30 Hz ERG amplitude based on the logged NGS-based final surface expression ratio. A multivariate model (ANOVA) was also used which adjusted for age at baseline visit, as younger subjects have higher baseline ERG amplitudes. For comparison to computationally-derived datasets, ΔΔG values (Rakoczy, 2011) and pathogenicity predictions from the Envision dataset (Uniprot P08100) (Gray, 2018) were tested as predictors of clinical severity as well.

## Results

### Example of inconclusive genetic testing results including a rare *RHO* VUS

This study hypothesizes that functional studies can help interpret DNA variants of unknown significance found in patient samples. Figure 2 shows an example of a clinical situation in which an inconclusive genetic testing result, including a rare rhodopsin VUS, was obtained for a proband with retinitis pigmentosa. Of the 210 *RHO* variants identified for this study, 10 variants were from rare *RHO* VUS that were discovered in DNA samples from subjects with inherited retinal diseases (See Methods and Table S1). Figure 2A shows the fundus appearance of one of these 10 subjects (D00726) who was diagnosed with retinitis pigmentosa after a full clinical evaluation. In this Mendelian disorder, it is most likely that variant(s) in a single gene are truly pathogenic. Figure 2B shows the large number of VUS identified in genetic testing for this subject using panel-based sequencing of inherited retinal disease genes (GEDi test)(Consugar, 2015). The *RHO* P180T variant (row 3) is predicted to be pathogenic, but so are other VUS to varying degrees, two of which are coding variants in genes that can cause dominantly-inherited retinitis pigmentosa (SNRNP200 and RP1). An expert in ocular genetics may suspect that the *RHO* variant is the most likely to be pathogenic, but it is not conclusive. Therefore, additional functional data are needed.

**Figure 2.**
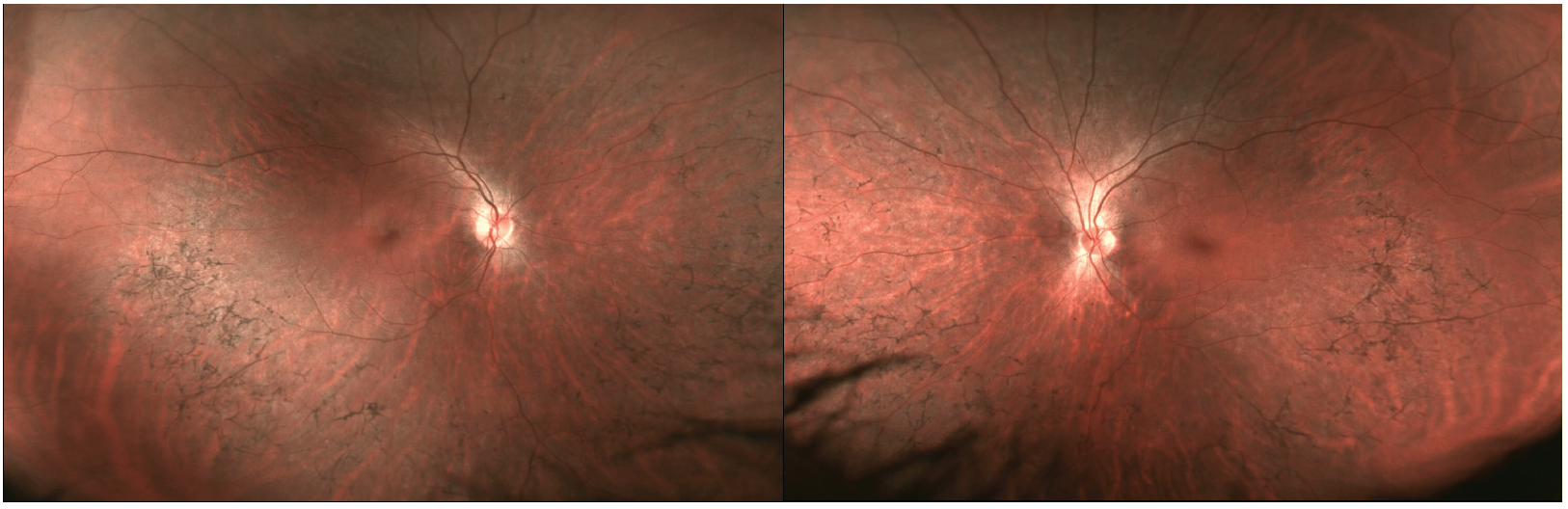
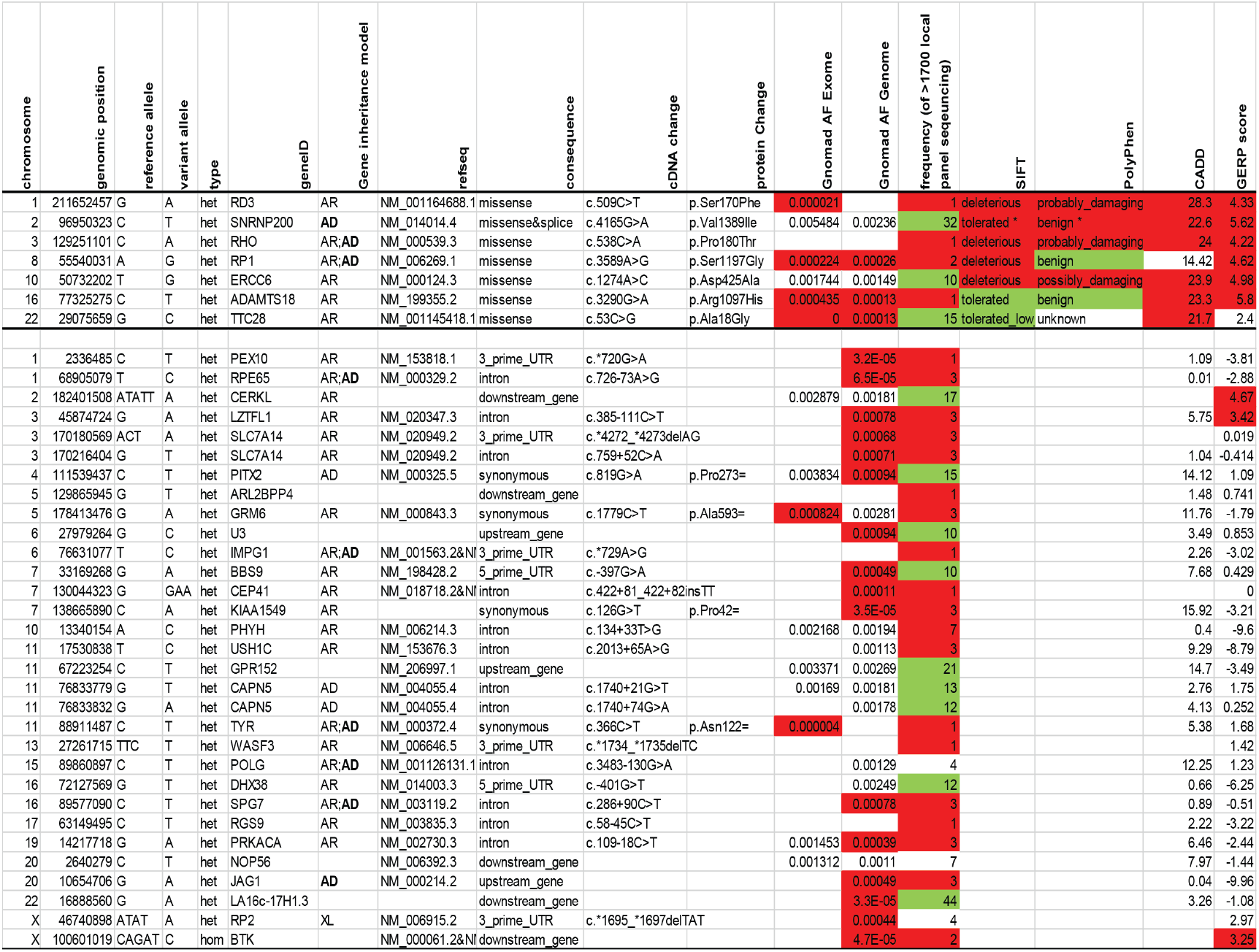
Example subject (ID# D00726) with multiple VUS. (A) Fundus photographs show midperipheral bone spicule pigmentation consistent with a diagnosis of RP. (B) DNA sequencing using the Gene Eye Disease panel revealed a large number of VUS, with annotations color coded (red= supporting pathogenicity; green= supporting benign). Variants above the horizontal line are coding variants. *splice site potentially broken, Human Splice Finder −34%.

### Flow cytometry assay validation

Rhodopsin variant plasmids (N=210) were created using site-directed mutagenesis and these were then cloned into an expression vector suitable for transfection into cultured cells. The next step (Figure 1) was to optimize a fluorescent assay to identify which variants have pathogenic surface expression levels. Based on the assay by McKeone et al (McKeone, 2014), Figure 3 shows that indirect immunofluorescence staining successfully distinguishes between wildtype RHO and known mutant (P23H) RHO. In this case, whereas the wildtype RHO was expressed well on the cell surface, the P23H mutant did not express on the cell surface, but rather was trapped inside the cell as expected. The results were validated using non-permeabilized cells to ensure that that only surface RHO was detected (Figures 3A and 3B); flow cytometry results corresponded to the immunofluorescence staining patterns(Fig. 3C). By comparing flow cytometry results from wildtype and P23H mutant RHO as positive and negative controls, respectively, the assay to separate wildtype from mutant RHO was highly reproducible (Z’=0.94, Fig. 3D) and therefore appropriate for use in a “high-throughput screening” context.

**Figure 3.**
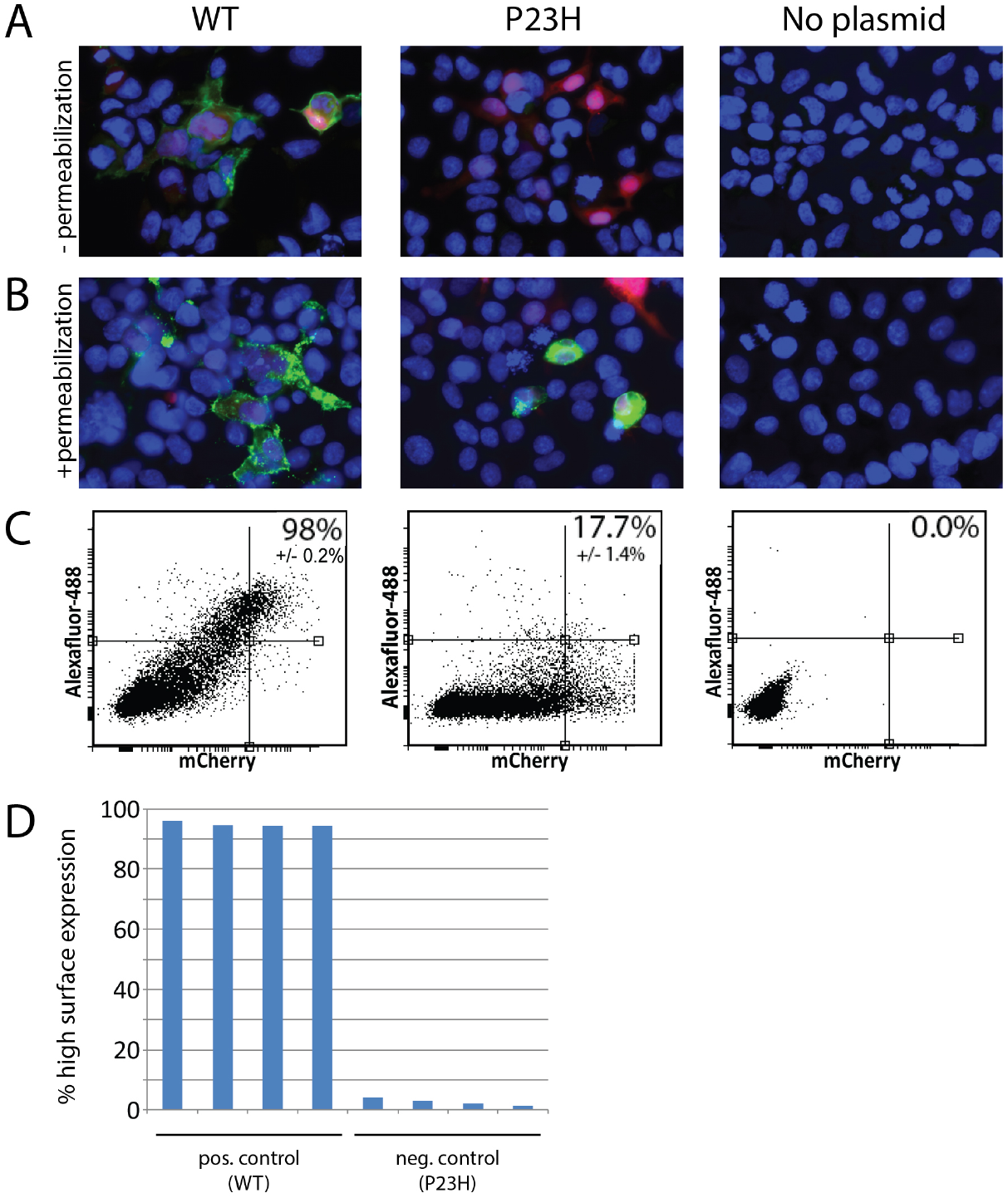
Unpooled assay validation. (A) Indirect immunofluorescence without permeabilization using Ret-P1/Alexafluor-488 antibodies shows strong RHO surface expression (green) using the wildtype plasmid (WT; left column), but not with a known mutant construct (P23H; middle column) or no plasmid control (right column). Red: mCherry transfection control. Blue: nuclei (Hoechst). (B) With permeabilization, mutant RHO is detectable inside the cell (middle column). (C) When the same antibodies were used in flow cytometry (without permeabilization), the percentage of RHO^+^/transfection^+^ cells (inset) correlates to the results obtained using immunofluorescence. (D) Replicate flow cytometry assays show high separation between positive and negative controls, and low noise, Z’= 0.94.

Each of the 210 variants in the library was then tested by flow cytometry using two methods for comparison: 1) the standard unpooled flow cytometry assay in which each variant is maintained in a separate tube and analyzed separately (Supplemental Figure 1), and 2) a pooled assay in which transfected cells are then pooled and FACS is used to sort RHO high-expressing versus low-expressing cells (Figure 4A). The entire cDNA within the transfected plasmid (inside the cell) serves as the “barcode” and allows deconvolution/quantitation of the level of each variant that is represented in the high and low pools. The residual plasmid is isolated from each cell pool, amplified by PCR and sequenced by NGS.

**Figure 4.**
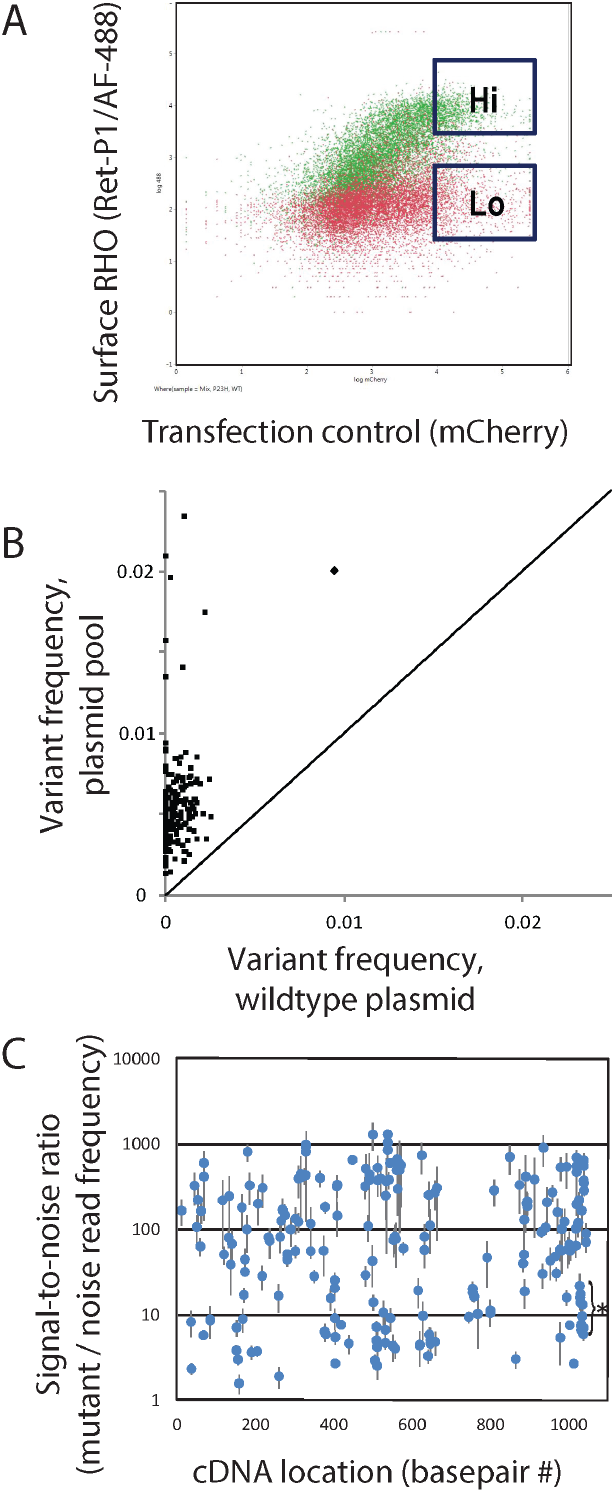
Pooled assay workflow and signal-to-noise ratio. (A) Gates used for FACS sorting cells with high or low rhodopsin surface expression, based on a positive control wildtype rhodopsin sample (green) and a negative control mutant sample (red). (B) Variant detection above background is demonstrated for each of 210 library variants, comparing sequencing results from of a pool of all variants (x-axis) to those from a wildtype *RHO* amplicon with no variants (y-axis). (C) Signal-to-noise ratio of the 210 variants as a function of location in the cDNA sequence. Error bars represent standard error of three independent replicates. Star = cluster of lower signal-to-noise probes-see text.

### Validation of string-based pooled variant detection and quantification

All 210 library variants were detected at a coverage-normalized frequency that was >1.5 of that observed by sequencing a wildtype *RHO* amplicon with no variants (Figure 4B). Two hundred and seven of the 209 variants were seen at >2x. One variant which was the used in the positive control sample during flow cytometry (diamond) showed overrepresentation, possibly due to cross-contamination at the flow cytometry stage; this was minimized in future iterations. Next, intra-sample quality control was performed by evaluating the level of the “noise probes” as described in the Methods. Customized computational filtering of sequencing quality scores was needed to obtain high signal-to-noise data. Figure 4C shows the signal-to-noise (S-N) ratio as a function of location in the cDNA. All variants were detected with a S-N ratio >1.5x, and with 208 of 210 detected at >2x (Figure 4C).

The starred cluster of points (Figure 4C) with a lower, but acceptable, S-N ratio is from variants located near a sequencing error-prone sequence context that is GC rich with a poly-C repeat (GGCCCCGGCC). There are a large number of reported variants in this region (Table S1), and in the 3’ end of the coding region in general, forming a “hot spot” of reported variants. In contrast, there are no reported variants between bp 665-744.

### Validation and quality control of the final NGS-based surface expression ratio

The final surface expression ratio was high when cells with a particular variant are primarily found by flow cytometry within the “Hi” gate (Figure 4A), with good surface expression, similar to wildtype RHO. The final ratio was low when cells with a particular variant are primarily found in the “Lo” gate, similar to known RHO mutants which do not express well on the cell surface. The characteristics of this ratio were evaluated both by comparing it to the standard unpooled flow cytometry assay and by its analytical and biological variability.

Figure 5 demonstrates that there was very good concordance between the standard unpooled assay and the NGS-based pooled assay, with an overall r^2^ of 0.92. The pooled assay drastically reduced the number of samples (per biological replicate) that had to be examined by flow cytometry; Unpooled: 210 samples analyzed plus three controls. Pooled: two samples sorted, plus three controls. For the minor discrepancies/outliers in the intermediate range of the assays, it is not known which assay was more accurate. The red and pink areas contain variants with ‘very low’ and ‘low’ expression levels, respectively, consistent with those variants being pathogenic mutations. While hard cutoffs of such regions have no exact biological meaning, for practical purposes the regions were defined by 40% and 80% surface expression on the x-axis, and the corresponding pooled assay ratios can be compared with positive and negative controls (Figure 6).

**Figure 5.**
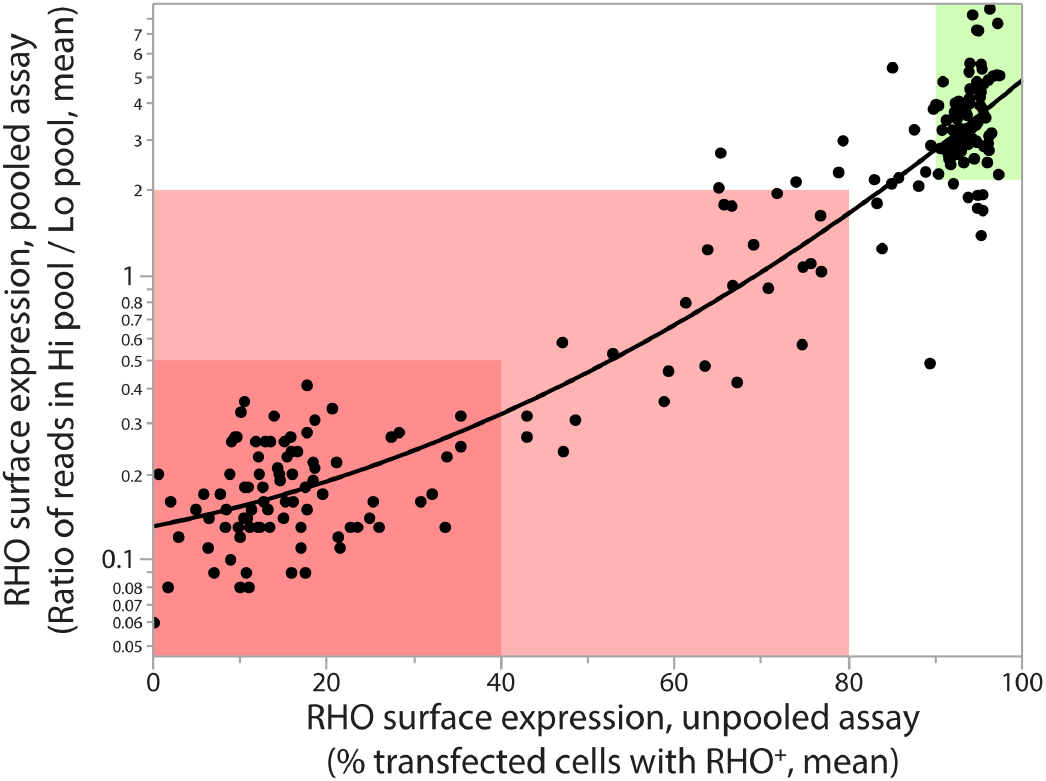
Concordance between unpooled and pooled assays. The unpooled RHO surface expression flow cytometry assay performed on individual tubes (x-axis) showed highly correlated results (quadratic fit; r^2^ =0.92) to the NGS-based, pooled assay (y-axis, log scale). Each point represents one variant. Red=very low expression; pink=low expression; green=high expression.

**Figure 6.**
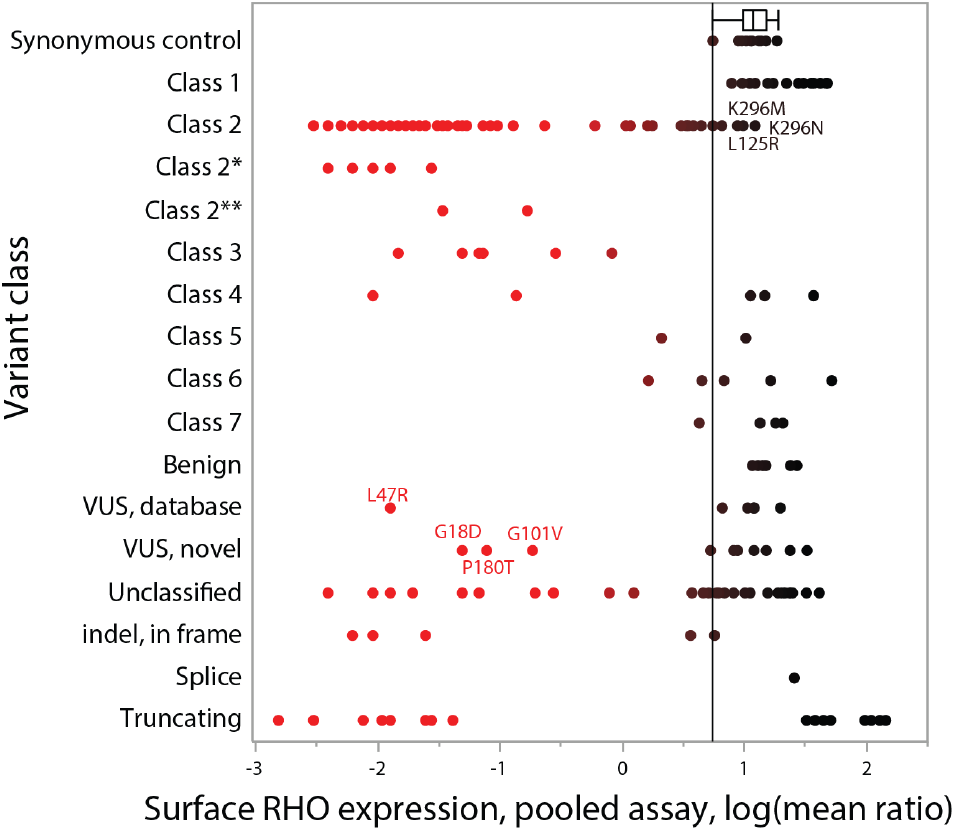
Pooled assay results by variant class. RHO surface expression results are plotted by previously-described biochemical classes. The color gradient (red-black) is based on the x-axis value, with pathogenic levels of expression to the left, and wildtype levels to the right. A vertical line is drawn at the lower bound of the box-and-whiskers plot, which shows the interquartile range and upper and lower data point values of the synonymous controls. Class 2*, following Athanasiou et al, might behave as class 4 after 11-cis-retinal rescue and Class 2** might behave as class 2 on overexpression, but class 4 in vivo. Labeled points are examples of variants previously considered class 2 mutants that unexpectedly show intermediate or high expression levels (upper) or VUS that were previously unclassified that are now likely to be class 2, 3, or 4 mutations (lower).

To evaluate variability, the pooled assay was performed on DNA from three separate biological replicates performed in different months. The resulting ratios showed high correlations between replicate experiments (r^2^=0.86-0.88; see Supplemental Figure 2). Of note, the data described above was derived from DNA amplification of the residual plasmid in the transfected cell. Ratios derived from RNA from the same sorted cells were much more variable, and while they showed similar trends, were less consistent (data not shown).

Next, the pooled assay results were graphed by the variant class derived from preexisting literature and database annotations (Figure 6). The reported biochemical classes were obtained from the literature (Mendes, 2005; Rakoczy, 2011; Athanasiou, 2018). (Additional historical categorizations have also been published (Sung, 1993; Krebs, 2010), as well as classifications that integrate pharmacologic response (Behnen, 2018).) The synonymous controls (top row)demonstrated high surface expression, as expected. Most of the known class 2 mutants exhibited very low surface expression as expected, but a minority (19%) expressed at levels that were unexpectedly high (L125R, E150K), intermediate (A164V, G51V, F52Y, F56Y, T58M, K296M, K296N) or moderately low (G51R, T58R, L88P, G109R, C167R, S186W, M207R, M216K, K296E). Class 3 variants showed intermediate levels. Class 4 variants showed low and high levels (see Discussion). Class 5-7 variants and benign variants show intermediate to high levels. Labeled variants that were previously considered VUS (L47R, G18D, G101V, and P180T) have now been demonstrated to express at pathogenic levels. Note that the variants with high expression levels are not necessarily all benign, as the assay does not detect all classes of mutations.

Next, six variants with new or unexpected findings were selected for further validation using immunofluorescence with confocal microscopy. A164V and G109R are class 2 mutants according to Rakoczy and Athanasiou(Rakoczy, 2011; Athanasiou, 2018), but unexpectedly showed intermediate or relatively preserved surface RHO staining in all three assays-standard and pooled flow cytometry (Table 1) as well as confocal microscopy (Figure 7). The following variants were originally of unknown pathogenicity: L47R (literature) and G18D, G101V, P180T (internal VUS). All four of these variants showed low or very low surface expression in the standard and pooled flow cytometry assays and were further validated to have low surface expression in the confocal immunofluorescence assay (Figure 7).

**Figure 7.**
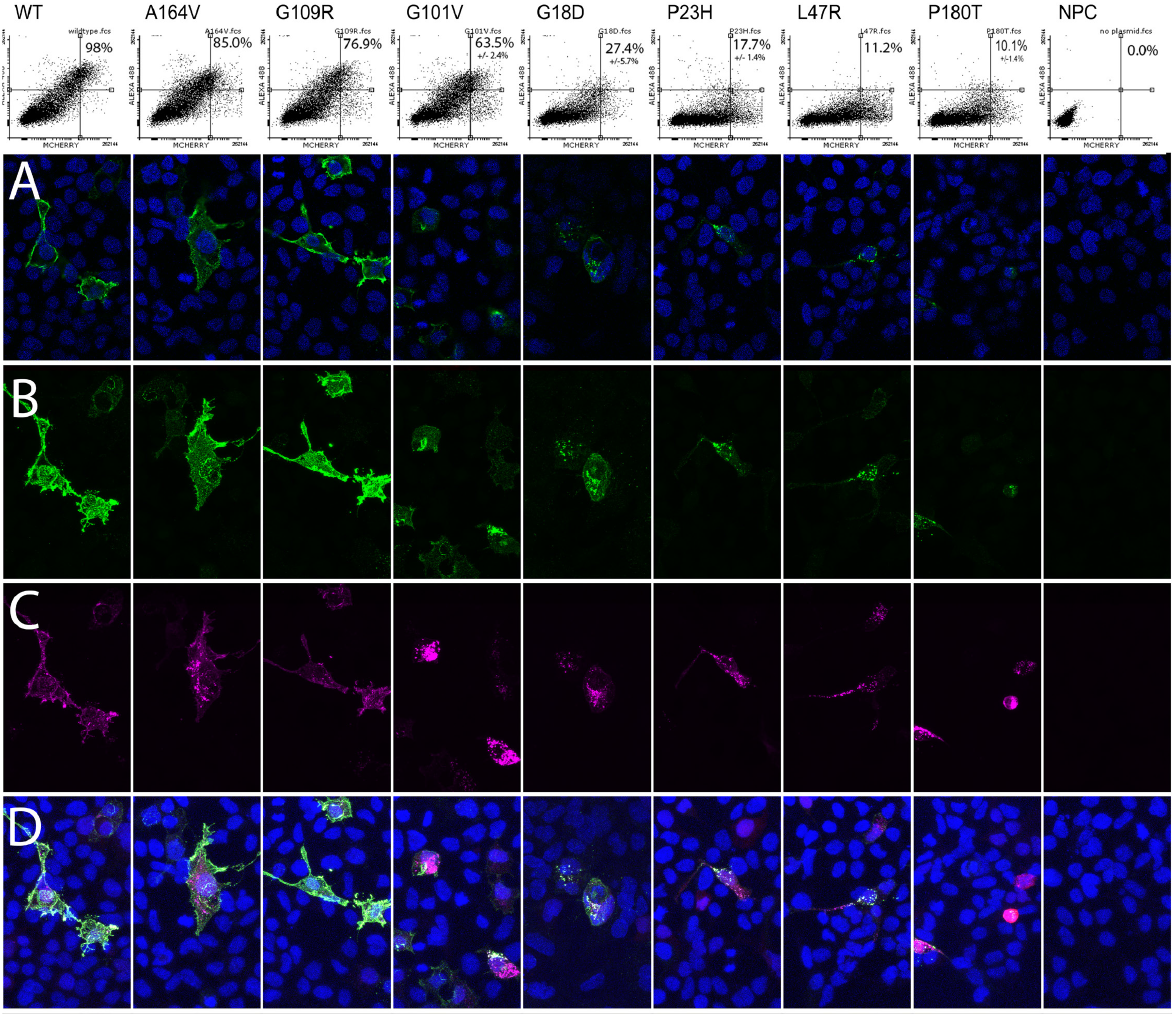
Confocal immunofluorescence of selected mutants validates flow cytometry results. (A) Each column represents a transfection with a particular labeled variant, arranged from highest (left) to lowest (right) RHO expression (WT=wildtype). Top row: Flow cytometry results showing surface RHO staining (Y axis) and transfection marker (X axis), with percent of transfected cells with high surface RHO as inset. Confocal staining: Green = RHO cell surface staining, Purple = total RHO staining after cell permeabilization, blue = nuclear stain. Row A: single-slice confocal images demonstrate cell membrane staining for the left-most three variants but fainter, punctate perinuclear staining for the other variants. A similar pattern of decreasing RHO intensity is seen in maximum projection images of (B) surface RHO and (C) total RHO. Row D shows a composite of all channels.

**Table 1.**
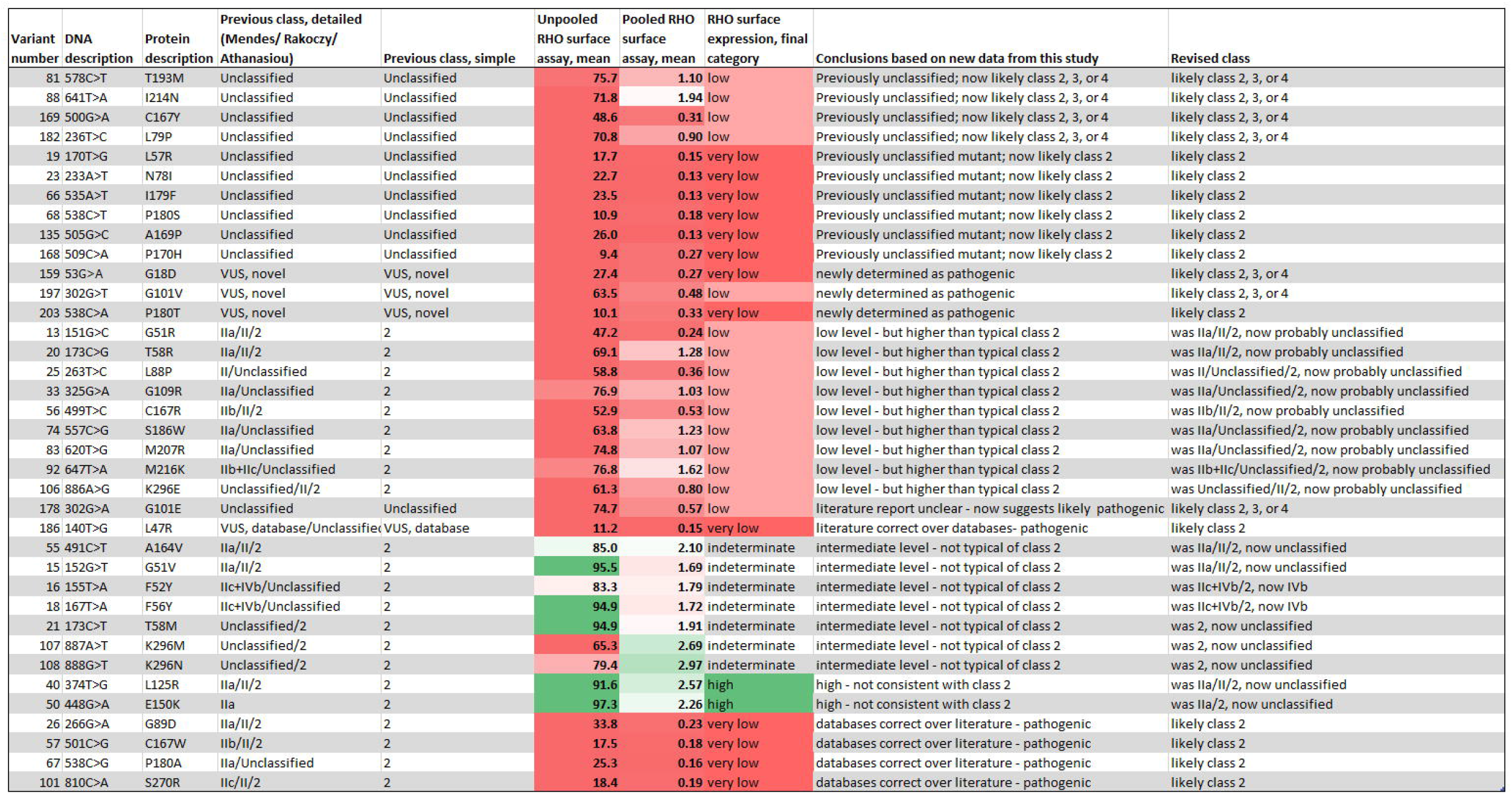
New findings based on RHO surface expression assay results. Previously-reported RHO mutation categories (“Previous class, detailed”) were revised (“Revised Class”) based on RHO surface expression results. Color coded gradients show pathogenic levels (red) and wildtype levels (green). For standardized variant descriptions (HGVS format), add prefixes NM_000539.3:c. for DNA and RHO_v001:p. or NP_000530.1:p. for protein descriptions.

### Probands with a rare *RHO* VUS

Ten probands that had genetic testing through our departmental genetic testing service had a novel *RHO* VUS, as listed in Table 2. These represent nine unique VUS (as G18D was found in two probands). (Since the start of this project, the p.R147C variant has been added to the HGMD database as a pathogenic variant.) Of the nine VUS, three (33%) showed pathogenic surface expression levels. The variants that were pathogenic by the functional assay were not uniformly predicted to be damaging based on computational predictions alone (Table 2). Both of the subjects with the G18D mutation and the one subject with the G101V mutation had a phenotype of pericentral retinitis pigmentosa, as described in further detail (Comander, 2017). The subject shown in Figure 2 who had retinitis pigmentosa and a P180T *RHO* variant now has been demonstrated to have a pathogenic *RHO* mutation.

**Table 2.**
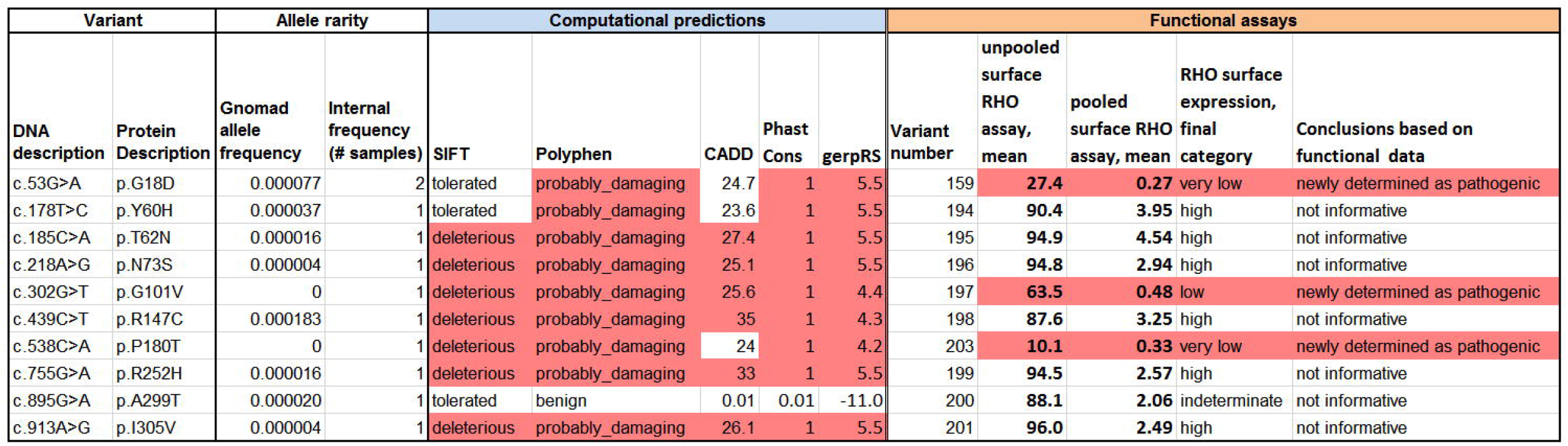
Comparison of information from computational methods and function assays to solve patient pedigrees. Among nine rare VUS that were identified from our genetic testing service, three variants had measured pathogenic expression levels. Color coding for computational methods are based on arbitrary cutoffs for SIFT (Ng, 2003), PolyPhen (Adzhubei, 2013), CADD scores (Kircher, 2014), PhastCons (Siepel, 2005), and GerpRS (Cooper, 2005). Color coding for functional assays is based on the RHO surface expression final category.

### Predicting clinical disease severity data from surface expression levels

Among known class 2 mutants, the level of surface RHO expression measured in vitro can predicted the clinical disease severity of class 2 mutants, with an observed increase of 0.67 of Ln ERG amplitude for each log of surface expression ratio (p=0.0008; Figure 8). A multivariate model that takes age-at-baseline visit into account gives similar results, with a 0.59 increase of ln ERG amplitude for each log of surface expression ratio (p=0.016). In contrast, ΔΔG values from Rakoczy et al(Rakoczy, 2011), representing the computationally predicted misfolding propensity, did not predict ERG amplitudes alone, or in a combined model with age or age and NGS ratio (all p>0.05). Similarly, computational pathogenicity predictions from the Envision dataset(Gray, 2018) did not correlate with clinical severity, with or without restriction to class 2 mutants (p>0.05).

**Figure 8.**
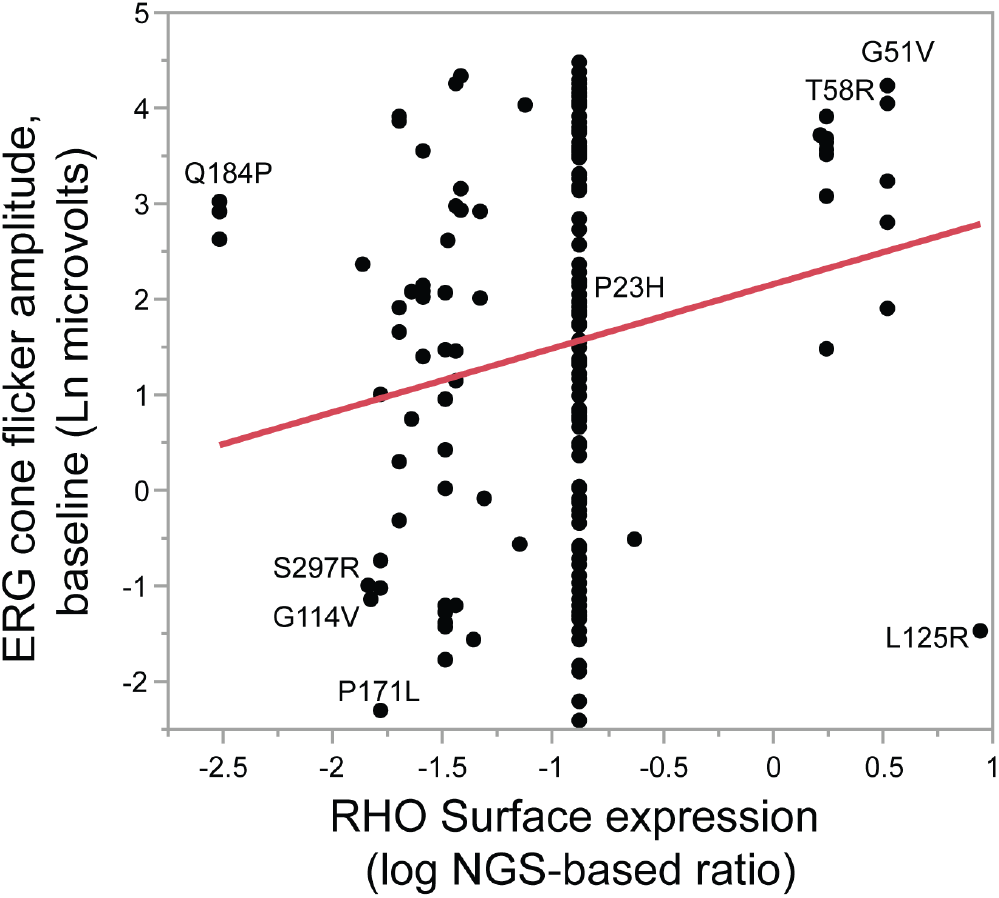
Correlating *RHO* surface expression and clinical disease severity. Among known class 2 RHO mutants, increasing amounts of RHO surface expression correlated with a milder clinical phenotype, as represented by the baseline cone flicker ERG response amplitude (p=0.012). Each point represents one patient.

## Discussion

This study demonstrates the feasibility of a higher-throughput cell-based assay for the functional characterization of VUS in IRDs, compared to the standard approach of using a cell-based assay for each variant individually (Sung, 1993; Li, 1998; Chuang, 2004; Chen, 2011; Toledo, 2011; Davies, 2012; Hollingsworth, 2013; Liu, 2013; McKeone, 2014; Yamasaki, 2014). A pooled, multiplexed assay for rhodopsin variants can efficiently identify class 2, 3, or 4a mutants. This assay was used to identify pathogenic variants within a group of VUS, provisionally solving 3 pedigrees of typical or pericentral retinitis pigmentosa, including the proband in Figure 2.

The barcode-free, pooled, NGS-based assay for evaluating RHO surface expression was highly reproducible and quantitative, with good agreement with the standard unpooled assay. This was achieved by optimizing several segments of the schema in Figure 1, including extensive library validation, with sequencing on both DNA strands after mutagenesis and after it was recombined into the final expression vector; optimization of fixation and staining conditions to maintain high discrimination between positive and negative controls, while also using a fixative that preserved DNA integrity for subsequent PCR; and quality filtering of NGS basecalls to maintain signal-to-noise in variant quantification.

The data in Tables 1 and S1 provide new information about previously-uncharacterized variants, identify apparently misclassified variants, and confirm pathogenicity of known mutations. Of 80 known class 2 mutants, 71 (89%) were confirmed to show pathogenically low levels of RHO surface expression. However, 9 (11%) class 2 mutants unexpectedly showed high or intermediate levels, especially those found in transmembrane helix I. 8 of 66 (12%) mutants in other mutation classes also showed pathogenic RHO surface expression levels, especially in class 3 and 4 mutants, which tended to have an intermediate phenotype. 14 of 33 (42%) unclassified pathogenic variants showed pathogenic expression levels, which are likely class 2, 3, or 4 mutants.

For example, *RHO* A292E does not cause RP but instead causes the milder disease congenital stationary night blindness. *RHO* A292E is known to be constitutively active in activating transduction without a chromophore (class 6) (Dryja, 1993). Computation predictions based on misfolding propensity (ΔΔG)(Rakoczy, 2011) led to the conclusion that this mutant should also be grouped in class 2 (IIa), but the relatively normal actual expression level of this variant indicates that it should be categorized as class 6 only. Conversely, T193M was predicted to fold correctly (Rakoczy, 2011), but in this study T193M is not expressed well on the cell surface. It is not known if this defect is due to misfolding or a hypothesized defect in dimerization (Rakoczy, 2011).

This expanded dataset also provides more confidence in variant interpretation based on smaller numbers of variants or among different experimental systems. For example, D190N showed low expression levels in this study, in contrast to smaller studies showing moderate expression using a GFP fusion tag (McKeone, 2014) or normal localization using a bovine sequence backbone (Liu, 2013).

Some special variant classes deserve further discussion. Even though RHO surface expression in HEK293 cells has been widely used to demonstrate pathogenicity (Sung, 1993; Li, 1998; Chuang, 2004; Chen, 2011; Toledo, 2011; Davies, 2012; Hollingsworth, 2013; Liu, 2013; McKeone, 2014; Yamasaki, 2014), recessive mutations do not show any special features in this assay, and are not distinguished from the dominant mutations. The recessive mutations are not clustered in a certain segment of the gene. However, because the number of known recessive mutants is small (6.6%; 12 of 181 according to Table S1), it is much more likely that any particular low-expressed mutation is dominant. Therefore, this study provides good evidence that the “very low” and “low” expressing variants are pathogenic mutations, but cannot conclusively state that they are *dominant* pathogenic mutations. (Thus, the three families mentioned above are “provisionally” solved.) Our initial experiments mixing together a known dominant mutation with a wildtype construct showed no effect on trafficking of the wildtype protein in this system (Supplemental Figure 3). Modifications of the cell type used or of expression levels may be able to reveal such dominant negative or gain of function effects to distinguish dominant pathogenic mutations from recessive pathogenic mutations. In a different system using the SK-N-SH cell type, dominant negative effects of dominant mutants have been demonstrated (Mendes, 2008).

This assay was focused on identifying class 2 (misfolding) mutations, which is the most common class. Figure 6 shows that class 3 mutations (endocytosis) and some class 4 mutations (altered post-translational modifications and reduced stability) also displayed low expression, though not quite as low as class 2, on average. Interestingly, the older Rakoczy classification separates the class 4 mutants slightly better, with the class IVa (affecting a glycosylation site; T4K, T17M, N15S) all showing pathogenic surface expression levels.

Additional assays are needed to detect and distinguish other mutation classes. For example, to identify mutations that are rescued by retinal binding(Mendes, 2008; Krebs, 2010; Mendes, 2010; Behnen, 2018), the assay could be repeated in the presence of retinal (which was too toxic for use in the current transient transfection system; not shown). For the purpose of evaluating pathogenicity, general toxicity- or stress-based assays would be ideal. In this cell culture system, expression of P23H or T17M mutants had no effect on cell death as measured by membrane permeability dyes, on apoptosis as measured by annexin V labeling, or on ER stress as measured by the pCAX-XBP1delDBD-venus reporter(Iwawaki, 2004)(not shown). Other approaches to look for these more general effects might use different cell types or even *in vivo*. Thus, a panel of both general and specific assays is likely the best approach to identify all types of pathogenic mutations.

Truncating mutations (nonsense or frameshift) were expressed in the context of a cDNA without internal introns, after the synthetic intron in the CAG promoter. Because internal introns and their potential effects of nonsense mediated decay (Roman-Sanchez, 2016) were not included, the results for those mutants should be interpreted with caution. All truncating variants 5’ of a.a. 315 showed pathogenic expression levels, while all variants 3’ to a.a. 332, including 10 truncating variants, showed wildtype expression levels. These variants should preferably be retested in their full genomic context (Roman-Sanchez, 2016), which would also allow for testing of splice variants.

### Screening small versus large variant libraries

This library construction strategy differs from larger-scale screening strategies(Melnikov, 2014; Brenan, 2016; Gasperini, 2016) (thousands of variants) in that this library, with 210 variants, was created with widely-available techniques at lower technical complexity. The library itself was constructed using modified Quikchange mutagenesis reactions using inexpensive short oligos ordered in 96 well plates. (Synthetic DNA variant libraries may replace this step as prices continue to decrease.) Each variant plasmid was introduced into cells using a standard transfection protocol, without need for viral vectors. The disadvantage of this approach is that it does not scale well to larger sized libraries. Sequencing and reracking of individual clones, as well as the unpooled transfection, become more cumbersome with increasing number of variants and wells. However, for biological questions such as the one addressed in this study involving about 200 variants or less, this approach is feasible and has the above advantages.

A smaller library size also has advantages in achieving a quantitative readout. In a very large, barcode-free, pooled assay, NGS sequencing errors would set a lower boundary so that the rarest variants are harder to quantify, and therefore these assays would best be used to find highly enriched “hits”. In this study, because of the smaller number of variants, the quantitation of each variant could be maintained above the noise floor caused by sequencing errors.

Therefore, smaller gradations in expression level were reproducibly quantified over the entire dynamic range of the assay (Figure S2), allowing for observations such as the “intermediate” expression phenotype of the Class 3 and 4 mutations described above. More specifically, for the quantitation of each of 210 plasmids in a mixed pool, the frequency of an individual plasmid (ideally ~1/210 = ~0.5%) has to be greater than the background noise of sequencing errors--theoretically 0.1% on the Illumina MiSeq at Q=30, though sequence-context and cycle number dependent. With library inhomogeneity and context-dependent sequencing error rates, this ratio is not guaranteed to be maintained. Careful masking of low quality basecalls allowed for low frequency variant quantitation above the noise level (Methods and Figure 4B-C). For larger barcode-free libraries, maintaining accurate quantification at low levels in the presence of read errors would not be solved by simply increasing read depth; specialized wet lab assays to lower the actual sequencing error rate (e.g. (Schmitt, 2012)) could be considered. Alternatively, barcoded libraries designed to avoid barcode collisions in the presence of sequencing errors could be used.

### Predicting clinical disease severity data from surface expression levels

Among known class 2 mutants, the level of RHO surface expression measured in this study was better at predicting disease severity than computational folding predictions (Figure 8 and Results). These approaches are not mutually exclusive, however, and structural biology can be used as a complementary approach to experimental observations. For example, folding calculations can be refined based on empirical data and then extended to new mutants. For variants that express well on the cell surface or where the mechanism is different (e.g. class 7 dimerization mutations) other classes of bioinformatic predictors are needed.

The outliers in Figure 8 that do not follow the general trend of less severe disease with increasing surface expression can be informative as well. For example, Q184P shows low severity despite low expression, and L125R shows high severity despite high expression (Figure 8). These data suggests that these variants are likely not classic class 2 mutations and influence disease course through a different mechanism than simple misfolding.

The correlation between surface expression and disease severity can produce some clinically-useful estimates. For example, a typical class 2 mutation with a low surface expression ratio of −1.5 would have a predicted baseline ERG which is 1.34 natural log units lower than a mutation with a higher expression ratio of 0.5. At an average rate of progression (0.091 ln units per year(Berson, 2002)), this corresponds to an extra 15 years of vision. If outliers (e.g. Q184P, L125R) in the regression had been excluded, this estimated effect would be larger. These estimates are based on the average severity for a mutant. However, there is a large amount of variation between individual subjects, particularly notable for the P23H mutation which is common in our cohort; additional modifying factors are yet to be identified, whether genetic, environmental, or stochastic.

In summary, a functional genomics approach can be used to address the problem of VUS in inherited retinal diseases, which in general is currently one of the major bottlenecks in the diagnosis of human Mendelian diseases. Future studies may include generalizing these assays to more genes and more mutation types, as well as using more complex methods to screen larger numbers of variants.

## Supporting information

Table 1 word format

Table 2 word format

## Acknowledgments

We would like to acknowledge Mark Consugar, Mindy Kwong, the CCIB DNA Core Facility at Massachusetts General Hospital for assistance with sequencing and colony processing, and David Dombrowski in the Massachusetts General Hospital Flow Cytometry Core for flow cytometry and flow sorting.

We are grateful for funding from NEI K12 EY016335-10 (JC), the Foundation Fighting Blindness Enhanced Career Development Award (JC), the Research for Prevention of Blindness Career Development Award (JC), the Massachusetts Lions Foundation / Lions Eye Research Foundation, and NEI/NIH P30 EY014104.

---

**Supplemental Data 1. Detailed spreadsheet of all variant information and results**. See annotation within the file.

**Supplemental Figure 1. Detailed display of unpooled flow cytometry results**. Individual flow cytometry plots (N=315) were sorted by the average NGS-based expression ratio (pooled assay), which visually demonstrated the good correlation between the unpooled and pooled assays. X-axis: mCherry co-transfection positive control Y-axis: anti-rhodopsin/AF-488. Plots at the top of the figure have a high NGS ratio, reflecting more cells in the upper-right quadrant of the flow cytometry plot (diagonal cloud of points). Plots at the bottom of the Figure shows a low NGS ratio, and a corresponding lack of points in the upper-right quadrant (horizontal cloud of points). Labels: “VAR”=variant number. “T”= transfection batch number

**Supplemental Figure 2. Pooled assay reproducibility**. Correlation coefficients (top) and scatterplot matrices (bottom) show good reproducibility between each replicate of the pooled, NGS-based assay (“NGS log ratio replicate” numbers 1, 2, and 3). The unpooled assay (“FACS average % high”) also shows good correlation with the mean pooled value (“NGS log ratio average”) and with the individual pooled replicates.

**Supplemental Figure 3. Dominant negative effect not observed**. Compared to expression of wildtype RHO alone (left), co-expression of the known dominant mutant RHO-P23H (right) does not prevent wildtype RHO from reaching the cell surface. X-axis: mCherry transfection control. Y-axis: anti-rhodopsin/AF-488.

