## Supplemental Figure 1 for "Characterizing variants of unknown significance in rhodopsin: a functional genomics approach"

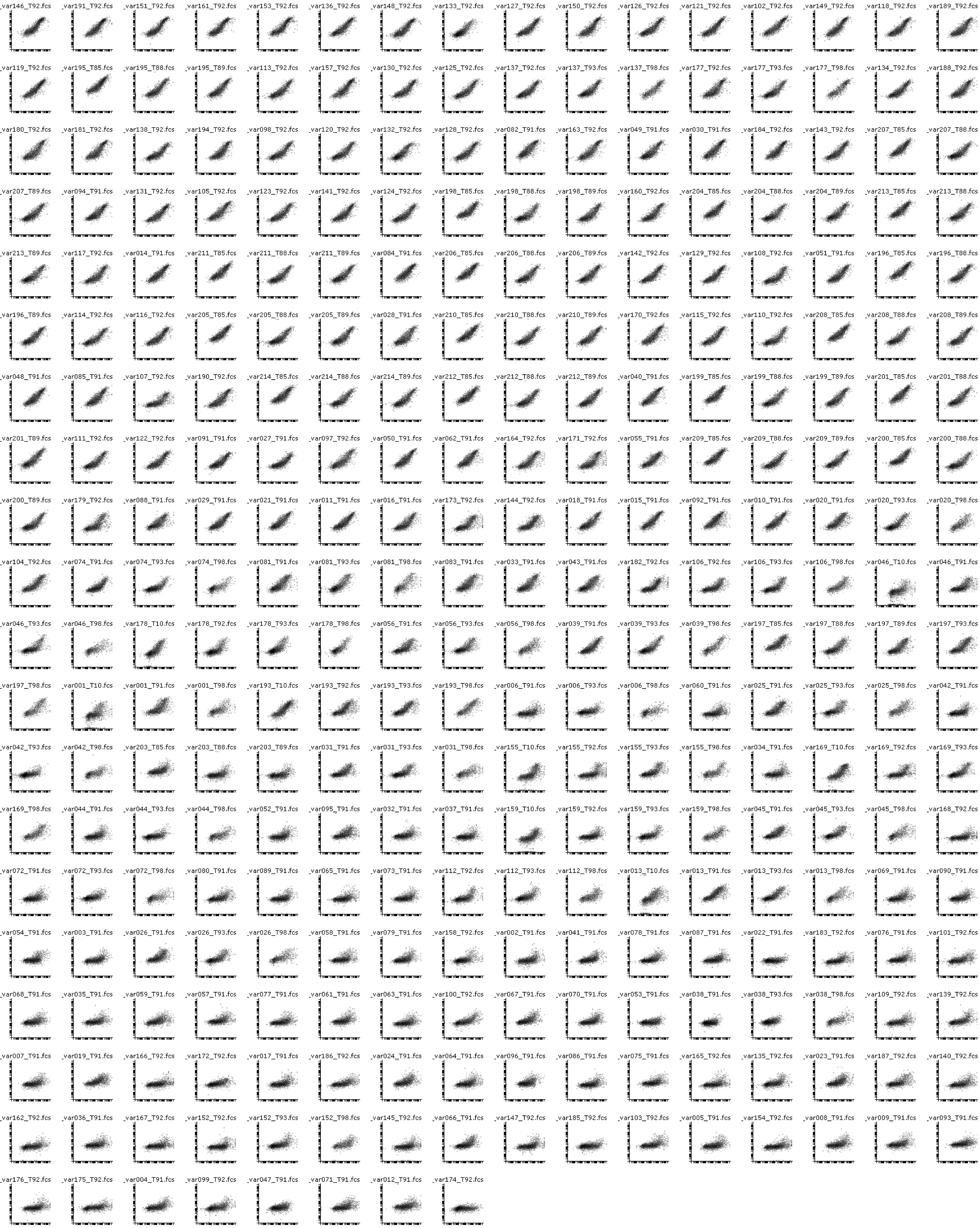


Detailed display of unpooled flow cytometry results. Individual flow cytometry plots (N=315) were sorted by the average NGS-based expression ratio (pooled assay), which visually demonstrated the good correlation between the unpooled and pooled assays.  X-axis: mCherry co-transfection positive control Y-axis: anti-rhodopsin/AF-488.  Plots at the top of the figure have a high NGS ratio, reflecting more cells in the upper-right quadrant of the flow cytometry plot (diagonal cloud of points).  Plots at the bottom of the Figure shows a low NGS ratio, and a corresponding lack of points in the upper-right quadrant (horizontal cloud of points). Labels: "VAR"=variant number. "T"= transfection batch number
