## Supplemental Figure 2 for "Characterizing variants of unknown significance in rhodopsin: a functional genomics approach"

### Correlations

| | FACS average %high | NGS log ratio average | NGS log ratio replicate 1 | NGS log ratio replicate 2 | NGS log ratio replicate 3 |
|---|---|---|---|---|---|
| FACS average %high | 1.0000 | 0.9492 | 0.9281 | 0.8833 | 0.9066 |
| NGS log ratio average | 0.9492 | 1.0000 | 0.9733 | 0.9413 | 0.9526 |
| NGS log ratio replicate 1 | 0.9281 | 0.9733 | 1.0000 | 0.8757 | 0.8838 |
| NGS log ratio replicate 2 | 0.8833 | 0.9413 | 0.8757 | 1.0000 | 0.8617 |
| NGS log ratio replicate 3 | 0.9066 | 0.9526 | 0.8838 | 0.8617 | 1.0000 |

### Scatterplot Matrix

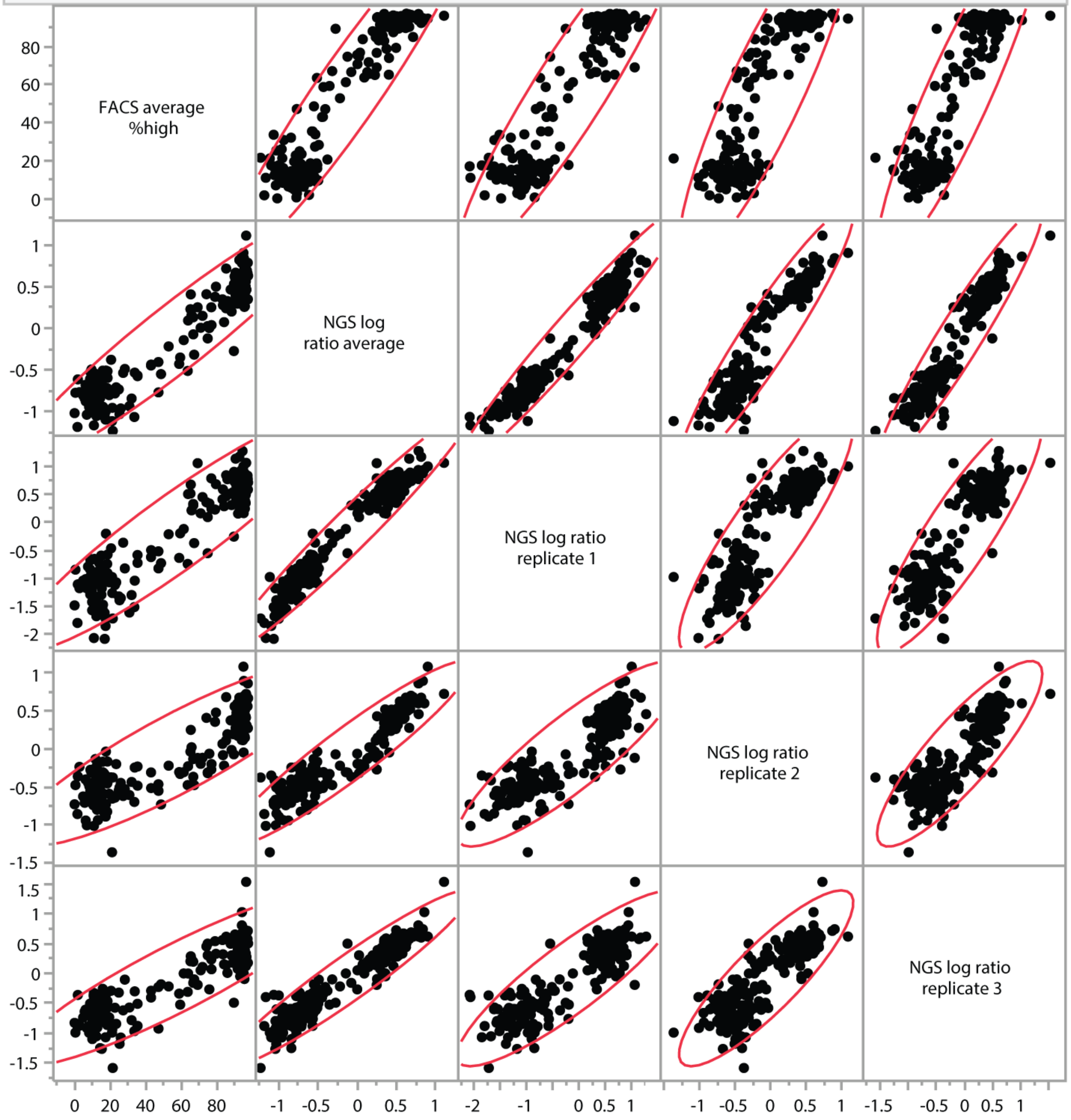


Pooled assay reproducibility. Correlation coefficients (top) and scatterplot matrices (bottom) show good reproducibility between each replicate of the pooled, NGS-based assay (“NGS log ratio replicate” numbers 1, 2, and 3). The unpooled assay (“FACS average % high”) also shows good correlation with the mean pooled value (“NGS log ratio average”) and with the individual pooled replicates.
