## Supplemental Figure 3 for "Characterizing variants of unknown significance in rhodopsin: a functional genomics approach"

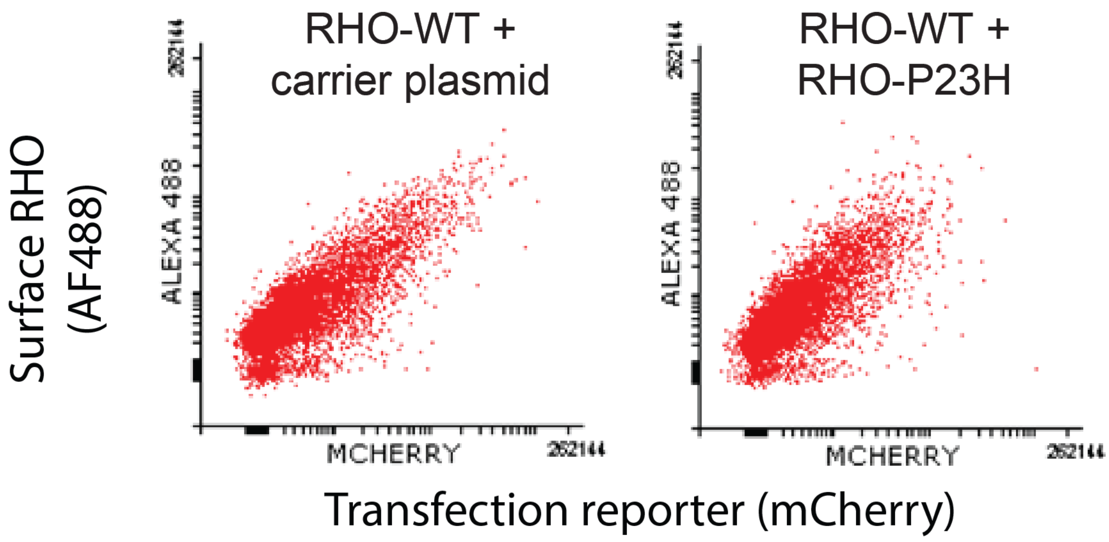


Dominant negative effect not observed. Compared to expression of wildtype RHO alone (left), co-expression of the known dominant mutant RHO-P23H (right) does not prevent wildtype RHO from reaching the cell surface. X-axis: mCherry transfection control. Y-axis: anti-rhodopsin/AF-488.
