## Supplementary material for "Characterizing variants of unknown significance in rhodopsin: a functional genomics approach": Table 1 word format

| **Variant number** | **DNA description** | **Protein description** | **Previous class, detailed (Mendes/ Rakoczy/ Athanasiou)** | **Previous class, simple** | **Unpooled RHO surface assay, mean** | **Pooled RHO surface assay, mean** | **RHO surface expression, final category** | **Conclusions based on new data from this study** | **Revised class** |
| --- | --- | --- | --- | --- | --- | --- | --- | --- | --- |
| 81 | 578C>T | T193M | Unclassified | Unclassified | **75.7** | **1.10** | low | Previously unclassified; now likely class 2, 3, or 4 | likely class 2, 3, or 4 |
| 88 | 641T>A | I214N | Unclassified | Unclassified | **71.8** | **1.94** | low | Previously unclassified; now likely class 2, 3, or 4 | likely class 2, 3, or 4 |
| 169 | 500G>A | C167Y | Unclassified | Unclassified | **48.6** | **0.31** | low | Previously unclassified; now likely class 2, 3, or 4 | likely class 2, 3, or 4 |
| 182 | 236T>C | L79P | Unclassified | Unclassified | **70.8** | **0.90** | low | Previously unclassified; now likely class 2, 3, or 4 | likely class 2, 3, or 4 |
| 19 | 170T>G | L57R | Unclassified | Unclassified | **17.7** | **0.15** | very low | Previously unclassified mutant; now likely class 2 | likely class 2 |
| 23 | 233A>T | N78I | Unclassified | Unclassified | **22.7** | **0.13** | very low | Previously unclassified mutant; now likely class 2 | likely class 2 |
| 66 | 535A>T | I179F | Unclassified | Unclassified | **23.5** | **0.13** | very low | Previously unclassified mutant; now likely class 2 | likely class 2 |
| 68 | 538C>T | P180S | Unclassified | Unclassified | **10.9** | **0.18** | very low | Previously unclassified mutant; now likely class 2 | likely class 2 |
| 135 | 505G>C | A169P | Unclassified | Unclassified | **26.0** | **0.13** | very low | Previously unclassified mutant; now likely class 2 | likely class 2 |
| 168 | 509C>A | P170H | Unclassified | Unclassified | **9.4** | **0.27** | very low | Previously unclassified mutant; now likely class 2 | likely class 2 |
| 159 | 53G>A | G18D | VUS, novel | VUS, novel | **27.4** | **0.27** | very low | newly determined as pathogenic | likely class 2, 3, or 4 |
| 197 | 302G>T | G101V | VUS, novel | VUS, novel | **63.5** | **0.48** | low | newly determined as pathogenic | likely class 2, 3, or 4 |
| 203 | 538C>A | P180T | VUS, novel | VUS, novel | **10.1** | **0.33** | very low | newly determined as pathogenic | likely class 2 |
| 13 | 151G>C | G51R | IIa/II/2 | 2 | **47.2** | **0.24** | low | low level - but higher than typical class 2 | was IIa/II/2, now probably unclassified |
| 20 | 173C>G | T58R | IIa/II/2 | 2 | **69.1** | **1.28** | low | low level - but higher than typical class 2 | was IIa/II/2, now probably unclassified |
| 25 | 263T>C | L88P | II/Unclassified | 2 | **58.8** | **0.36** | low | low level - but higher than typical class 2 | was II/Unclassified/2, now probably unclassified |
| 33 | 325G>A | G109R | IIa/Unclassified | 2 | **76.9** | **1.03** | low | low level - but higher than typical class 2 | was IIa/Unclassified/2, now probably unclassified |
| 56 | 499T>C | C167R | IIb/II/2 | 2 | **52.9** | **0.53** | low | low level - but higher than typical class 2 | was IIb/II/2, now probably unclassified |
| 74 | 557C>G | S186W | IIa/Unclassified | 2 | **63.8** | **1.23** | low | low level - but higher than typical class 2 | was IIa/Unclassified/2, now probably unclassified |
| 83 | 620T>G | M207R | IIa/Unclassified | 2 | **74.8** | **1.07** | low | low level - but higher than typical class 2 | was IIa/Unclassified/2, now probably unclassified |
| 92 | 647T>A | M216K | IIb+IIc/Unclassified | 2 | **76.8** | **1.62** | low | low level - but higher than typical class 2 | was IIb+IIc/Unclassified/2, now probably unclassified |
| 106 | 886A>G | K296E | Unclassified/II/2 | 2 | **61.3** | **0.80** | low | low level - but higher than typical class 2 | was Unclassified/II/2, now probably unclassified |
| 178 | 302G>A | G101E | Unclassified | Unclassified | **74.7** | **0.57** | low | literature report unclear - now suggests likely pathogenic | likely class 2, 3, or 4 |
| 186 | 140T>G | L47R | VUS, database/Unclassified | VUS, database | **11.2** | **0.15** | very low | literature correct over databases- pathogenic | likely class 2 |
| 55 | 491C>T | A164V | IIa/II/2 | 2 | **85.0** | **2.10** | indeterminate | intermediate level - not typical of class 2 | was IIa/II/2, now unclassified |
| 15 | 152G>T | G51V | IIa/II/2 | 2 | **95.5** | **1.69** | indeterminate | intermediate level - not typical of class 2 | was IIa/II/2, now unclassified |
| 16 | 155T>A | F52Y | IIc+IVb/Unclassified | 2 | **83.3** | **1.79** | indeterminate | intermediate level - not typical of class 2 | was IIc+IVb/2, now IVb |
| 18 | 167T>A | F56Y | IIc+IVb/Unclassified | 2 | **94.9** | **1.72** | indeterminate | intermediate level - not typical of class 2 | was IIc+IVb/2, now IVb |
| 21 | 173C>T | T58M | Unclassified/2 | 2 | **94.9** | **1.91** | indeterminate | intermediate level - not typical of class 2 | was 2, now unclassified |
| 107 | 887A>T | K296M | Unclassified/2 | 2 | **65.3** | **2.69** | indeterminate | intermediate level - not typical of class 2 | was 2, now unclassified |
| 108 | 888G>T | K296N | Unclassified/2 | 2 | **79.4** | **2.97** | indeterminate | intermediate level - not typical of class 2 | was 2, now unclassified |
| 40 | 374T>G | L125R | IIa/II/2 | 2 | **91.6** | **2.57** | high | high - not consistent with class 2 | was IIa/II/2, now unclassified |
| 50 | 448G>A | E150K | IIa | 2 | **97.3** | **2.26** | high | high - not consistent with class 2 | was IIa/2, now unclassified |
| 26 | 266G>A | G89D | IIa/II/2 | 2 | **33.8** | **0.23** | very low | databases correct over literature - pathogenic | likely class 2 |
| 57 | 501C>G | C167W | IIb/II/2 | 2 | **17.5** | **0.18** | very low | databases correct over literature - pathogenic | likely class 2 |
| 67 | 538C>G | P180A | IIa/Unclassified | 2 | **25.3** | **0.16** | very low | databases correct over literature - pathogenic | likely class 2 |
| 101 | 810C>A | S270R | IIc/II/2 | 2 | **18.4** | **0.19** | very low | databases correct over literature - pathogenic | likely class 2 |

Table 1. Previously-reported *RHO* mutation categories (“Previous class, detailed”) were revised (“Revised Class”) based on RHO surface expression results. Color coded gradients show pathogenic levels (red) and wildtype levels (green). For standardized variant descriptions (HGVS format), add prefixes NM_000539.3:c. for DNA and RHO_v001:p. or NP_000530.1:p. for protein descriptions.
