## Supplementary material for "Characterizing variants of unknown significance in rhodopsin: a functional genomics approach": Table 2 word format

| **Variant** | | **Allele rarity** | | **Computational predictions** | | | | | **Functional assays** | | | | |
| --- | --- | --- | --- | --- | --- | --- | --- | --- | --- | --- | --- | --- | --- |
| **DNA description** | **Protein Description** | **Gnomad allele frequency** | **Internal frequency (# samples)** | **SIFT** | **Polyphen** | **CADD** | **PhastCons** | **gerpRS** | **Variant number** | **unpooled surface RHO assay, mean** | **pooled surface RHO assay, mean** | **RHO surface expression, final category** | **Conclusions based on functional data** |
| c.53G>A | p.G18D | 0.000077 | 2 | tolerated | probably_damaging | 24.7 | 1 | 5.5 | 159 | **27.4** | **0.27** | very low | newly determined as pathogenic |
| c.178T>C | p.Y60H | 0.000037 | 1 | tolerated | probably_damaging | 23.6 | 1 | 5.5 | 194 | **90.4** | **3.95** | high | not informative |
| c.185C>A | p.T62N | 0.000016 | 1 | deleterious | probably_damaging | 27.4 | 1 | 5.5 | 195 | **94.9** | **4.54** | high | not informative |
| c.218A>G | p.N73S | 0.000004 | 1 | deleterious | probably_damaging | 25.1 | 1 | 5.5 | 196 | **94.8** | **2.94** | high | not informative |
| c.302G>T | p.G101V | 0 | 1 | deleterious | probably_damaging | 25.6 | 1 | 4.4 | 197 | **63.5** | **0.48** | low | newly determined as pathogenic |
| c.439C>T | p.R147C | 0.000183 | 1 | deleterious | probably_damaging | 35 | 1 | 4.3 | 198 | **87.6** | **3.25** | high | not informative |
| c.538C>A | p.P180T | 0 | 1 | deleterious | probably_damaging | 24 | 1 | 4.2 | 203 | **10.1** | **0.33** | very low | newly determined as pathogenic |
| c.755G>A | p.R252H | 0.000016 | 1 | deleterious | probably_damaging | 33 | 1 | 5.5 | 199 | **94.5** | **2.57** | high | not informative |
| c.895G>A | p.A299T | 0.000020 | 1 | tolerated | benign | 0.01 | 0.01 | -11.0 | 200 | **88.1** | **2.06** | indeterminate | not informative |
| c.913A>G | p.I305V | 0.000004 | 1 | deleterious | probably_damaging | 26.1 | 1 | 5.5 | 201 | **96.0** | **2.49** | high | not informative |

Table 2. Nine rare variants of unknown significance identified from our genetic testing service, comparing information from computational methods[ref] (center columns) and function assays (right columns). Color coding for computational methods are based on abritrary cutoffs for CADD scores (25), PhastCons (1), and GerpRS (3). Color coding for Functional assays is based on the final category. Both subjects with the G18D mutation and the one subject with the G101V mutation had a phenotype of pericentral retinitis pigmentosa, as recently published [ref]. R147C was added to HGMD since the start of this project
